# Chitin soil amendment triggers systemic plant disease resistance through enhanced pattern-triggered immunity

**DOI:** 10.1101/2024.12.08.627391

**Authors:** Moffat Makechemu, Yukihisa Goto, Helen Zbinden, Victoria Widrig, Beat Keller, Cyril Zipfel

**Author notes:** Correspondance : Cyril Zipfel.

## Abstract

Chitin triggers localised and systemic plant immune responses, making it a promising treatment for sustainable disease resistance. However, the precise molecular mechanisms underlying chitin-induced systemic effects in plants remain unknown. In this study, we investigated the effects of soil amendment with crab chitin flakes (hereafter chitin) on pattern-triggered immunity (PTI) and systemic disease resistance in various plant species. We found that soil amendment with chitin potentiates PTI and disease resistance against the bacterial pathogen *Pseudomonas syringae* pv. tomato DC3000 in lettuce, tomato, and Arabidopsis as well as against the fungal pathogen *Blumeria graminis* causing powdery mildew in wheat. Using micrografting in Arabidopsis, we demonstrated that this systemic effect is dependent on active chitin perception in the roots. We also showed that induced systemic resistance (ISR) and pattern-recognition receptors (PRRs)/co-receptors, but not systemic acquired resistance (SAR), are involved in the systemic effects triggered by chitin soil amendment. This systemic effect correlated with the transcriptional up-regulation of key PTI components in distal leaves upon chitin soil amendment. Notably, chitin-triggered systemic immunity was independent of microbes present in soil or chitin flakes. Together, these findings contribute to a better understanding of chitin-triggered systemic immunity, from active chitin perception in roots to the potentiation of PTI in the leaves, ultimately priming plants to mount enhanced defense responses against pathogen attacks. Our study provides valuable insights into the molecular mechanisms of chitin soil amendment and resulting induced immunity, and highlights its potential use for sustainable crop protection strategies.

## Introduction

Plants face persistent challenges from various pathogens, such as bacteria and fungi, which can have a substantial impact on crop yield and quality (Reddy and Reddy, 2015; Singh et al., 2023). An important aspect of plant immunity is the detection of pathogen/microbe-associated molecular patterns (PAMPs/MAMPs) by pattern-recognition receptors (PRRs) on the plant cell surface, leading to pattern-triggered immunity (PTI) that restricts pathogen growth (Ngou et al., 2024). Hallmarks of PTI include among others, calcium influx, callose deposition, production of reactive oxygen species (ROS) and stomatal closure (Yu et al., 2017). Activation of such local defense responses can induce the production of long-distance defense signals to activate systemic immune responses in distal tissues (Sun and Zhang, 2021). Systemic acquired resistance (SAR) is a defense mechanism where localised leaf infection triggers signaling molecules, prompting distant leaves to activate immune responses (Vlot et al., 2021), while induced systemic resistance (ISR) is classically initiated by beneficial bacteria residing in the plant rhizosphere (Mauch-Mani et al., 2017; Pieterse et al., 2014).

Chitin is the second most abundant and important natural biopolymer after cellulose. It is a linear polysaccharide composed of repeating beta-1,4-linked 2-acetamido-2-deoxy-d-glucopyranose (GlcNAc) units, which serves as a prominent structural element in the cell walls of fungi and the exoskeletons of various organisms such as insects, nematodes, and crustaceans (*e.g.* crabs, crawfish, lobsters, and shrimps) (Liaqat & Eltem, 2018; Rinaudo, 2006). Chitin is recognized as a PAMP/MAMP, detected as non-self by host PRRs which activates PTI (Gong et al., 2020; Li et al., 2020; Shinya et al., 2015). Notably chitin can also have growth-promoting properties and induces various physiological responses in certain plant species, particularly under stress conditions (Turk, 2019; Zhao et al., 2019).

The mechanisms of chitin perception and signaling in plants were first reported in rice (*Oryza sativa*, *Os*), through the identification of CHITIN-ELICITOR BINDING PROTEIN (CEBiP), which contains three extracellular LysM motifs anchored to the plasma membrane via a glycosylphosphatidylinositol (GPI)-anchor (Gong et al., 2017, Gong et al., 2020; Kaku et al., 2006). In rice, chitin perception triggers the formation of a heterodimer complex between OsCEBiP and the LysM-receptor kinase (LysM-RK) CHITIN ELICITOR RECEPTOR KINASE 1 (OsCERK1). Thus, in rice, two LysM proteins, OsCEBiP and OsCERK1, are necessary for chitin perception and signaling (Hayafune et al., 2014; Shimizu et al., 2010). In *Arabidopsis thaliana* (*At*, hereafter, Arabidopsis), a homolog of OsCERK1 called AtCERK1/LYK1 plays a crucial role in chitin as well as bacterial peptidoglycan perception (Miya et al., 2007; Wan et al., 2008; Gimenez-Ibanez et al., 2009; Willmann et al., 2011; Wang et al., 2017;). Homodimers of AtCERK1/LYK1 directly bind long-chain chitin oligomers (Liu et al., 2012). However, other studies proposed that other LysM-RKs, such as AtLYK5 and AtLYK4, which possess inactive kinase domains, may serve as primary chitin receptors, with contrasting binding affinities to chitin and roles in chitin-induced signaling (Cao et al., 2014; Mittendorf et al., 2024). Of note, in the context of certain immune responses, such as plasmodesmata closure, CERK1 is dispensable, and LYK4 forms a complex with the CEBiP ortholog LYM2 (Cheval et al., 2020). In tomato (*Solanum lycopersicum*, *Sl*), a CERK1 ortholog, Bti9 (also named SlLYK13) is required for chitin signaling and full antibacterial immunity (Zeng et al., 2012). In wheat (*Triticum aestivum*, *Ta*), TaCEBiP, TaLYK5 and TaCERK1 were also recently shown to be required for chitin responses (Liu et al., 2023).

The agricultural applications of chitin have gained increasing attention in recent years (Li et al., 2020; Riseh et al., 2024). Chitin possesses good biocompatibility, biodegradability, nontoxicity and various biological activities and poses no known hazard to the environment (Sharp, 2013). Notably, treatment with chitin or its deacetylated derivative, chitosan, has been shown to improve resistance to diseases caused by fungi, oomycetes, bacteria, viruses, and nematodes in multiple plant species (including crops), consistent with the perception of chitin/chitosan as PAMP/MAMP by the plant immune system (Gong et al., 2020; Dave et al., 2021; Riseh et al., 2024). While most reported uses of chitin (or chitosan) in this context have been through ectopic plant treatments of above ground plant tissue, the addition of chitin to soil represents an interesting alternative approach. It was shown to confer resistance to soil-borne pathogens and pests, and to increase soil suppressiveness to such pathogens (Riseh et al., 2024). Furthermore, due to its chemical properties and effect on the soil microbial composition, chitin has great potential for improving soil quality, including in the context of continuous cropping agricultural practices (Debode et al., 2016; Fan et al., 2022). While the local defense responses induced by chitin are well-established, emerging evidence suggests that chitin can also elicit systemic effects, enhancing disease resistance in distant plant parts. For example, Takagi et al. (2022) demonstrated that soil amendment with chitin induces systemic disease resistance in rice, which is dependent on the chitin receptors OsCERK1 and OsCEBiP. Disruption of these genes impairs resistance against the fungal pathogen *Bipolaris oryzae* and affects the expression of chitin-responsive genes. These findings suggest that chitin triggers systemic resistance through a mechanism involving long-distance signaling (Debode et al., 2016; De Tender et al, 2019; Shamshina et al., 2020; Takagi et al., 2022). However, there is a notable gap in our understanding of the precise molecular mechanisms behind chitin-induced systemic effects in plants.

In this study, we investigated the underlying molecular mechanisms involved in chitin-induced systemic effects upon chitin soil amendment and their impact on plant defense signaling pathways. We employed genetic approaches, including the use of mutants lacking specific PRRs and co-receptors, ISR and SAR key regulators, to investigate the involvement of key signaling components mediating chitin-induced systemic effects. We utilized Arabidopsis as a model system and extended our investigations to economically important crops such as tomato, lettuce (*Lactuca sativa*, *Ls*), and wheat. We evaluated disease resistance against bacterial and fungal pathogens, as well as PTI immune readouts such as ROS production and calcium influx. The findings of this study make a substantial contribution to our comprehension of the molecular mechanisms that underlie chitin-induced systemic resistance and its potential application in crop protection.

## Results and Discussion

### Chitin soil amendment promotes plant disease resistance against pathogens

We first conducted disease resistance assays to evaluate the systemic effects of chitin soil amendment in various plant species against different pathogens. We infiltrated Arabidopsis, tomato, and lettuce leaves of plants grown on chitin-amended soil with the hemibiotrophic bacterial pathogen *Pseudomonas syringae* pv. *tomato* DC3000 (*Pst* DC3000) and inoculated wheat with the biotrophic fungus *Blumeria graminis* f. sp*. tritici* (*Bgt)* causing powdery mildew (hereafter Bg.tritici). We observed a significant reduction in *Pst* DC3000 growth in Arabidopsis, tomato and lettuce following chitin soil amendment (Fig. 1A-C). Additionally, wheat subjected to chitin soil amendment exhibited suppressed growth of Bg.tritici (Fig 1D). These findings confirm that chitin soil amendment enhances systemic disease resistance in multiple plant species against both bacterial and fungal pathogens, highlighting chitin’s broad-spectrum potential as a natural amendment for enhancing plant resistance. This aligns with previous research that has documented similar enhancements in resistance following chitin amendment (Ai et al., 2023; Debode et al., 2016; Takagi et al., 2022).

**Figure 1:**
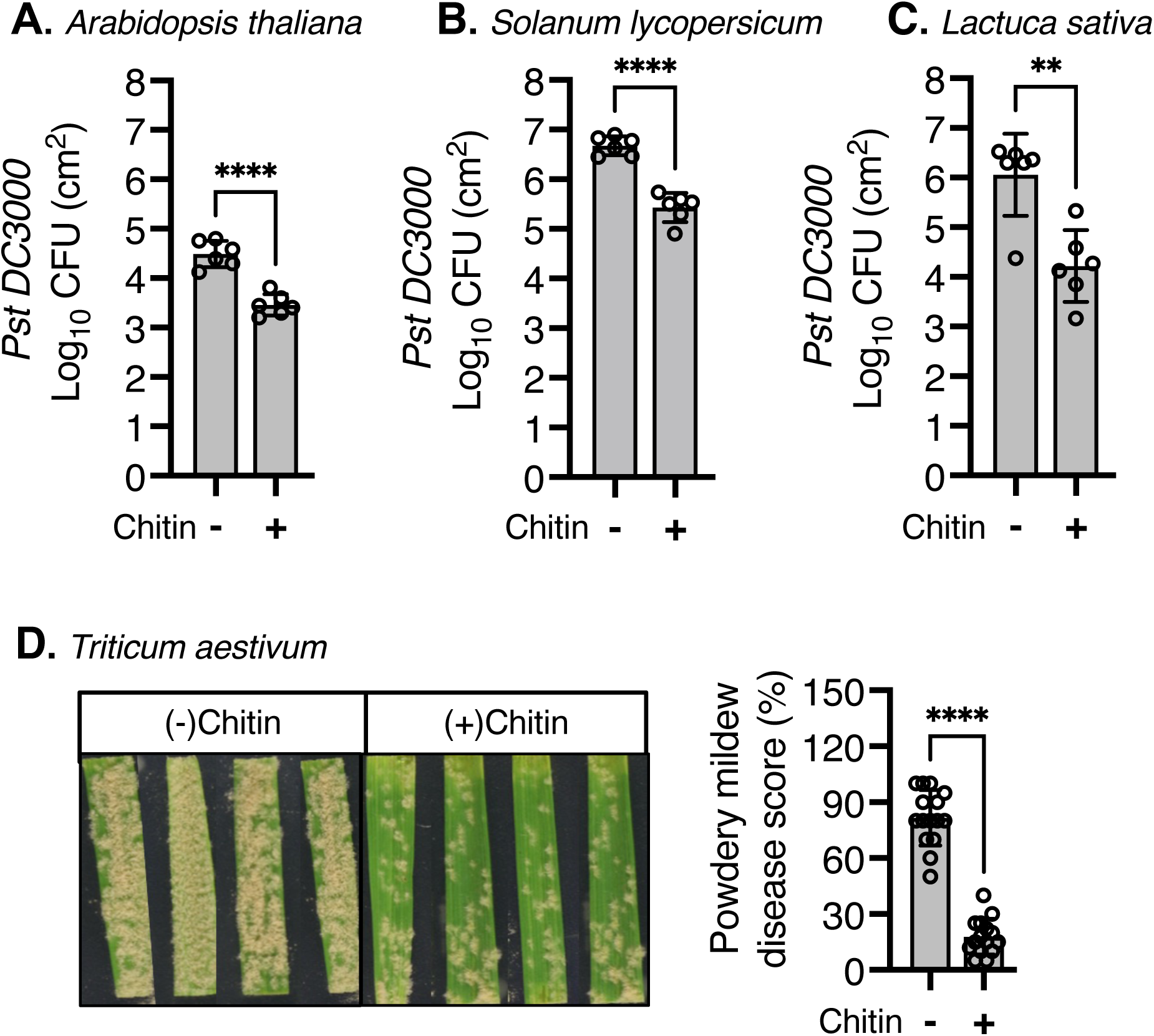
Chitin soil amendment suppresses growth of different leaf pathogens in different plant species. (A–C) Four-week-old *Arabidopsis thaliana* plants of Col-0 (A), tomato (*Solanum lycopersicum*) cv. Rio Grande (B), and lettuce (*Lactuca sativa*) cv. Salinas (C), grown in chitin-amended (+) or non-chitin (-) soil, were syringe-infiltrated with *Pst* DC3000 at a concentration of OD_600_=0.0002. Leaf disks were harvested two days post-infiltration, ground, and plated to quantify bacterial titers. Colony counts were performed two days post-incubation at 28 °C. Values represent mean ± standard error (SE, n=6). (D) Two-to three-cm-long leaf segments from 12-day-old wheat (*Triticum aestivum*) cv. Chinese Spring were challenged with *Blumeria graminis f. sp. tritici*, and powdery mildew symptoms were quantified nine days post-inoculation based on Kaur et al., (2008). Values are mean ± standard error (SE, n=12). Asterisks indicate a significant difference based on Student’s *t*-test (****, P ≤ 0.0001).

### Chitin perception is required for chitin-induced systemic effects

Chitin perception is mediated by the LysM-RK family (LYKs) across various plant species, demonstrating its evolutionary conservation (Gong et al., 2020; Li et al., 2020; Wang et al., 2017; Liu et al., 2023; Zeng et al., 2012).To explore the role of perception in chitin-induced systemic effects, we examined *Atcerk1* mutants in Arabidopsis and *bti9*/*lyk13* mutants in tomato, which exhibit compromised chitin perception. Induced resistance assays on these mutants revealed a significant reduction in systemic effects against *Pst* DC3000 (Fig. 2A-B). Extending our studies to lettuce, we generated *Lscerk1* mutants impaired in chitin perception using CRISPR-Cas9 to target the *LOC111905667* gene, an ortholog of *AtCERK1* (Fig. S1-S4). Induced resistance assay showed reduced chitin-induced systemic effects against *Pst* DC3000 in *Lscerk1* mutants upon soil amendment with chitin (Fig. 2C). The decrease in systemic induced resistance in multiple chitin receptor mutants across diverse plant species highlights the critical role of chitin perception in initiating systemic immune responses.

**Figure 2:**
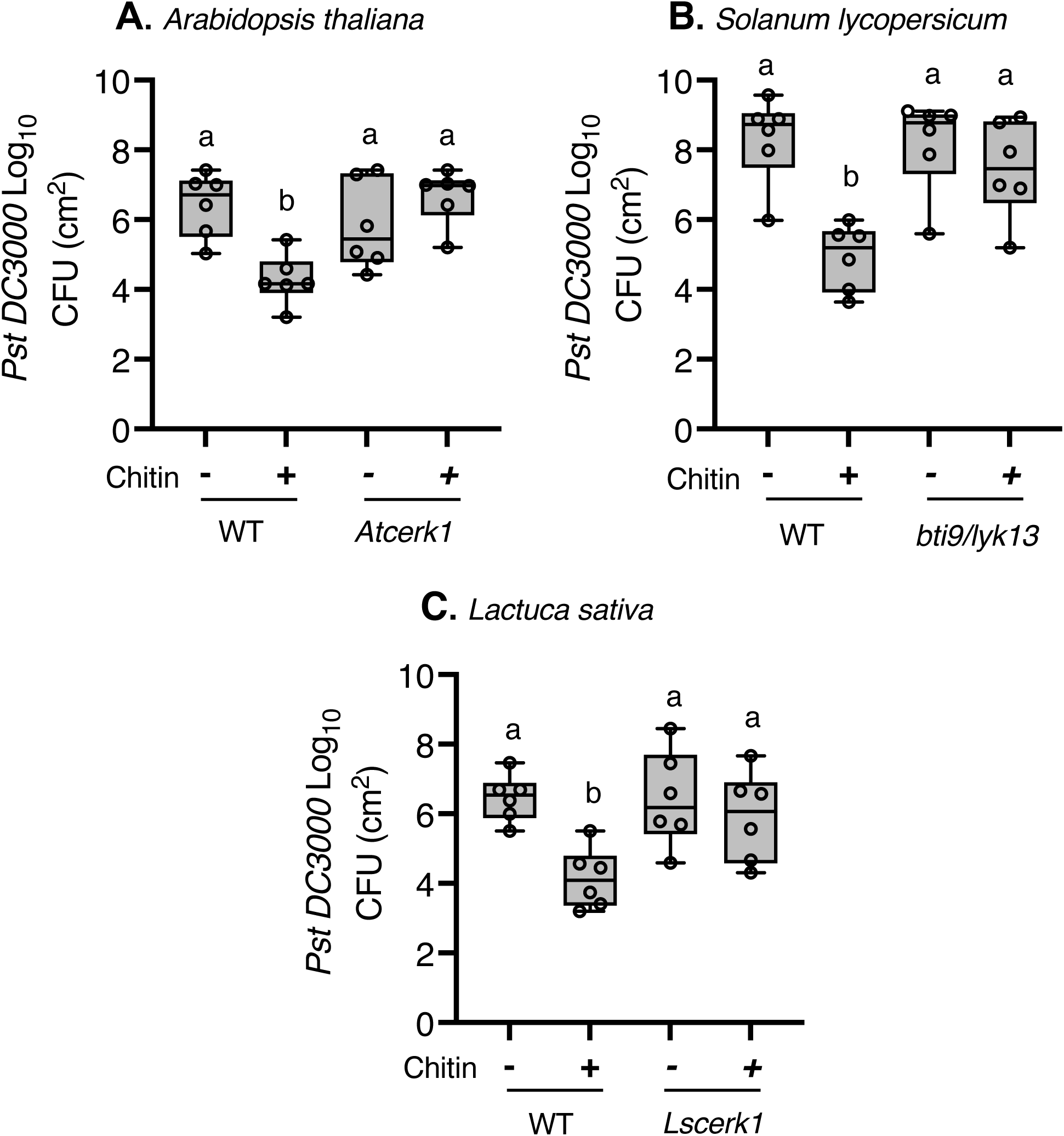
*cerk1* mutants show compromised chitin-induced systemic resistance in plants. (A–C) Four-week-old plants of *Arabidopsis thaliana,* Col-0 (WT) and *cerk1* (A), tomato (*Solanum lycopersicum*) Rio Grande WT and *bti9/lyk13* (B), and lettuce (*Lactuca sativa*) Salinas WT and *Lscerk1* (C), grown in chitin-amended (+) or non-chitin (-) soil, were syringe-infiltrated with *Pst* DC3000 at a concentration of OD_600_=0.0002. Leaf disks were harvested two days post-infiltration, ground, and plated to quantify bacterial titers. Values represent mean ± SE (n=6).

### Chitin perception in the roots initiates chitin-induced systemic effects

Micrografting has been instrumental in elucidating systemic signaling mechanisms in plants, enabling significant insights into processes such as shoot-to-root and root-to-shoot signaling, as well as flowering, nutrient response, and gene silencing (Tsutsui et al., 2020; Vanderstraeten et al., 2022; Tsutsui and Notaguchi, 2017; Molnar et al., 2010; Notaguchi et al., 2008; Pant et al., 2008; Tabata et al., 2014; Takahashi et al., 2018). To further explore the functional relevant tissues of chitin perception, we used micrografting of Arabidopsis *Atcerk1* mutants and wild-type Col-0 to create chimeric plants. These included combinations of *Atcerk1* (rootstock) grafted to Col-0 (scion), and *vice versa*, with Col-0-to-Col-0 as control. Induced resistance assays revealed that chimeric plants with *Atcerk1* as rootstock showed compromised chitin-induced disease resistance against *Pst* DC3000 (Fig. 3). In contrast, chimeric plants with Col-0 as rootstock maintained resistance levels similar to control plants (Fig. 3). This demonstrates the critical role of chitin perception in roots in initiating initiating induced resistance to leaf infection.

**Figure 3:**
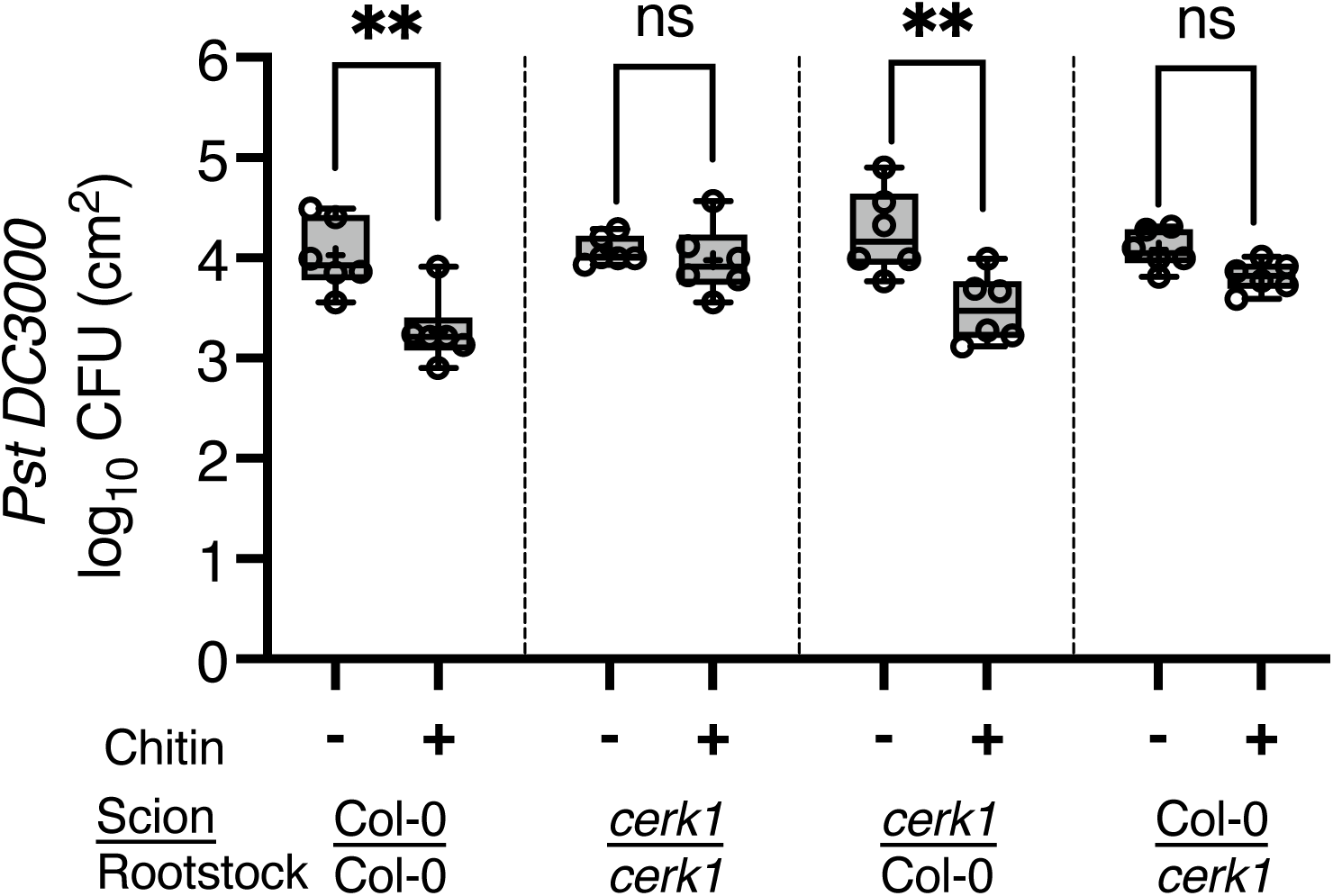
*cerk1* root stock chimeric Arabidopsis plants are impaired in chitin-induced systemic disease resistance. Five-week-old *Arabidopsis thaliana* chimeric plants, all grown in chitin (+) vs non-chitin (-) amended soil were syringe-infiltrated with *Pst* DC3000 at a concentration of OD_600_=0.0002. Harvested leaf disks were ground and plated for colony counting to quantify bacterial titers two days post infiltration. The mean ± standard error of the mean (SEM) was calculated from three independent biological replicates. Statistical significance is denoted by asterisks (*), where ** represents p<0.01.

### Chitin soil amendment potentiates flg22- and elf18-induced ROS production and calcium influx

After observing the crucial role of local chitin perception in roots for systemic effects and the suppression of pathogens in leaves, we investigated the involvement of PTI in leaves upon chitin-induced systemic effects. We focused on two commonly used PTI readouts namely ROS production and calcium influx. We conducted experiments in Arabidopsis, tomato, and lettuce upon soil amendment with chitin. In Arabidopsis, we elicited the immune responses using flg22 and elf18 (Fig. 4A), while only flg22 was used in tomato (Fig. 4B) and lettuce (Fig. 4C), given that elf18 perception is Brassicaceae-specific. We found that chitin soil amendment significantly enhanced ROS production in leaves of all plant species tested, compared to plants without chitin soil amendment (Fig. 4A-C). Similar results were observed in Arabidopsis lines expressing Aequorin (AEQ), a calcium-sensitive reporter, in which chitin soil amendment potentiated calcium influx (Fig. 4D). These findings indicate that chitin soil amendment can potentiate PTI outputs in leaves.

**Figure 4:**
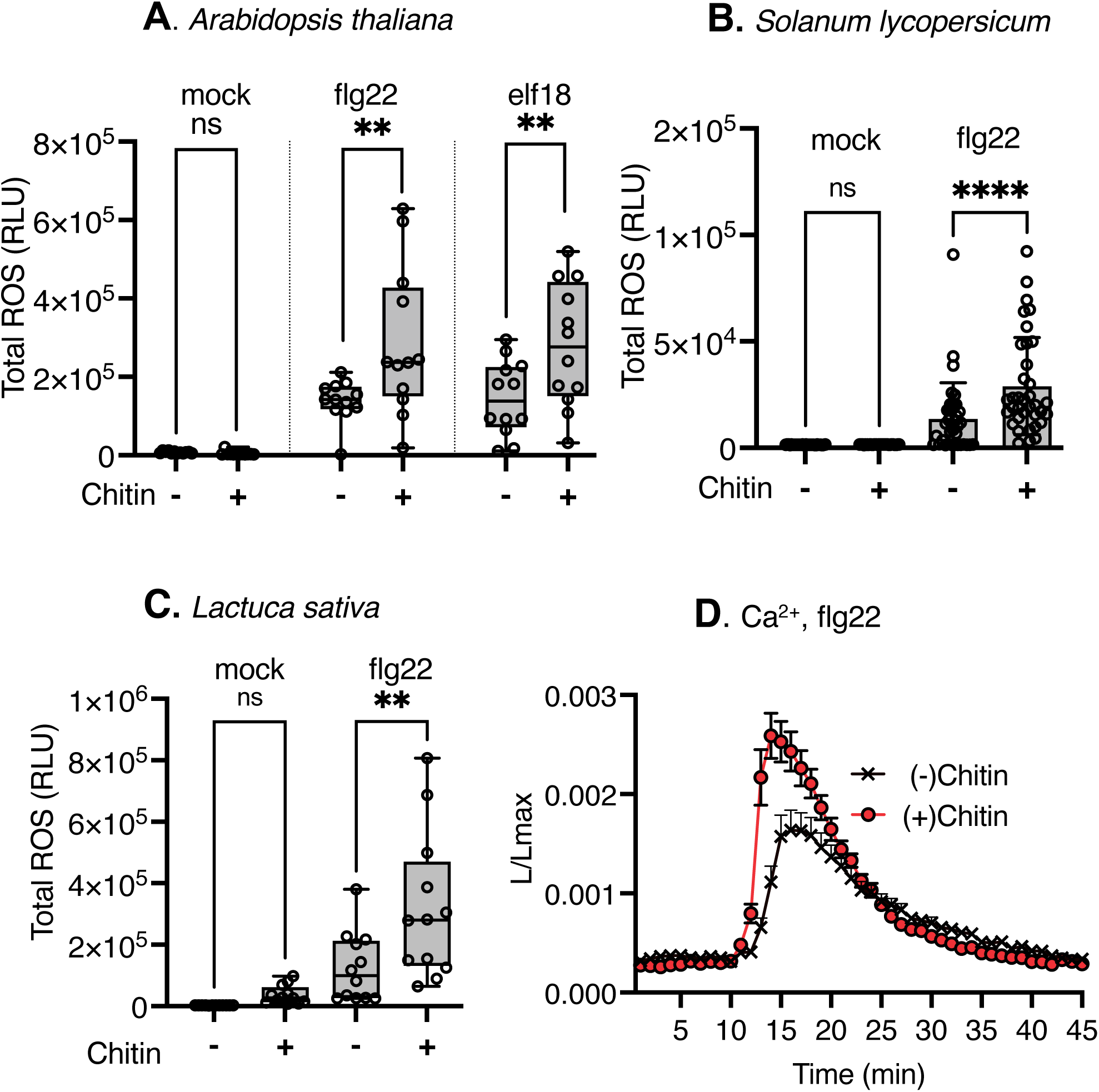
Chitin soil amendment potentiates ROS production and calcium influx in leaves. A–C) Leaf disks from 4-week-old plants of *Arabidopsis thaliana* Col-0 (A), tomato (*Solanum lycopersicum*) cv. Rio Grande (B), and lettuce (*Lactuca sativa)* cv. Salinas (C), grown in chitin-amended (+) or non-chitin (-) soil, were treated with 100 nM elf18 (A only), flg22, or mock. Total ROS relative light units (RLU) were quantified 42 minutes post-treatment. Values represent mean ± SE (n=12 for A; n=36 for B and C). Asterisks indicate significant differences based on Student’s t-test (**, P ≤ 0.01; ***, P ≤ 0.001). (D) Leaf disks from 4-week-old Col-0-AEQ plants grown in chitin-amended (+) or non-chitin (-) soil were treated with 100 nM flg22 or mock. ROS RLU was normalized to pre-treatment values (0–10 min), and total RLU was quantified 45 minutes post-treatment. Values represent mean ± SE (n=12).

### Chitin-induced systemic effects are linked to common PRR/co-receptors in Arabidopsis

Given the critical role of PTI in plant defense and the observed potentiation of PTI responses upon chitin soil amendment in our study, we aimed to dissect the contributions of well-characterized PRRs/co-receptors to chitin-mediated systemic effects in Arabidopsis, making use of extensive availability of genetic resources in this model plant. We tested several mutants of PRRs/co-receptors by conducting induced resistance assays on 5-week-old Arabidopsis plants grown with or without chitin soil amendment. Notably, most of the tested PRR/co-receptor mutants showed reduced disease resistance in chitin-treated leaves compared to untreated ones, whereas this chitin-induced effect remained intact in the wild-type Col-0 (Fig. 5).These findings, based on a genetic approach, suggest a correlation between chitin perception in roots and the involvement of PRRs/co-receptors in leaves, as evidenced by the increased susceptibility of PRR/co-receptor mutants to *Pst* DC3000, even in the presence of chitin

**Figure 5:**
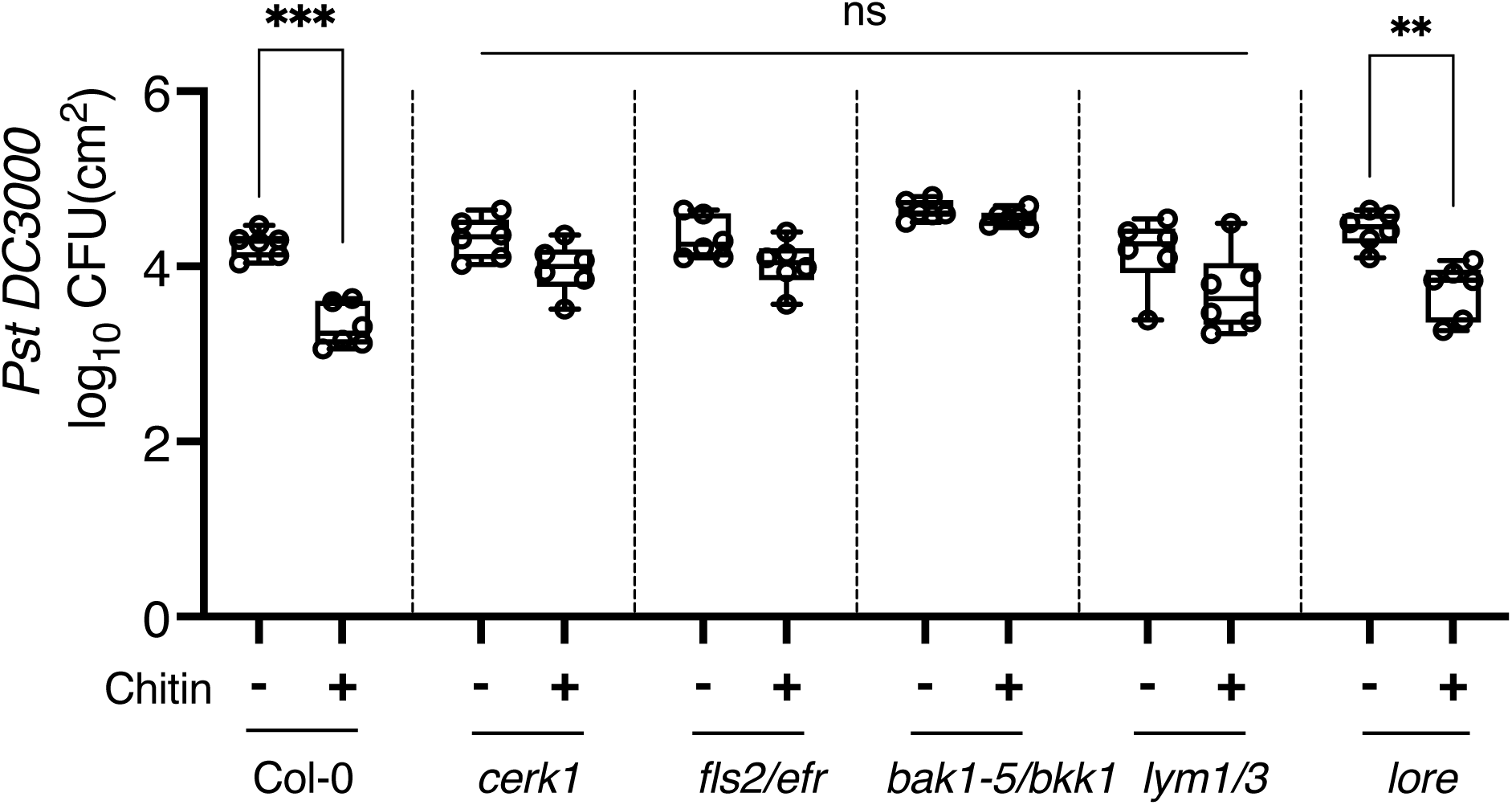
Chitin-induced systemic effects are linked to well-charactised PRRs or co-receptors in Arabidopsis. Four-week-old *Arabidopsis thaliana* Col-0 and mutants of PRRs/co-receptors (*bak1-5/bkk1*, *fls2/efr*, *cerk1*, *lym1/3* and *lore*) grown in chitin (+) vs non-chitin (-) amended soil were syringe-infiltrated with *Pst* DC3000 at a concentration of OD_600_=0.0002. Harvested leaf disks were ground and plated for colony counting to quantify bacterial titers two days post-infiltration. Values are mean ± SE, (n=6). Asterisks indicate a significant difference based on Student’s t-test (**, P≤0.01; ***, P≤0.001).

### Chitin-induced systemic effects involves ISR but not SAR

To understand the mechanism by which chitin perception in roots translates to systemic defense responses in distal plant tissues, we focused on the interplay between the well-established plant defense pathways of SAR and ISR. SAR is traditionally understood to function in leaf-to-leaf defense signaling, while ISR is known to convey signals from roots to leaves, likely offering broad-spectrum protection against a variety of pathogens (Mauch-Mani et al., 2017; Pieterse et al., 2014; Vlot et al., 2021) We examined Arabidopsis mutants associated with these pathways to investigate the link between roots and leaves in mediating the systemic effects of chitin. The mutants were specifically chosen for their roles in SAR (*fmo1* and *ald1*) and ISR (*dde2/ein2/pad4/sid2, npr1/ein2/jar1,myc2, bglu42*) (Vlot et al., 2021). This approach allowed us to see whether the observed systemic immune responses, such as enhanced disease resistance and flg22-induced ROS production, could be attributed to one or both pathways. Our findings revealed that ISR mutants showed compromised chitin-induced disease resistance (Fig. 6) and reduced flg22-induced ROS production (Fig. S5), indicating that the ISR pathway likely plays a major role in mediating chitin-induced systemic disease resistance. In contrast, SAR mutants (*fmo1*, *ald1*) did not show similar impairment (Fig. 6), further suggesting that ISR, rather than SAR, is the primary pathway responsible for the systemic effects of chitin.

**Figure 6:**
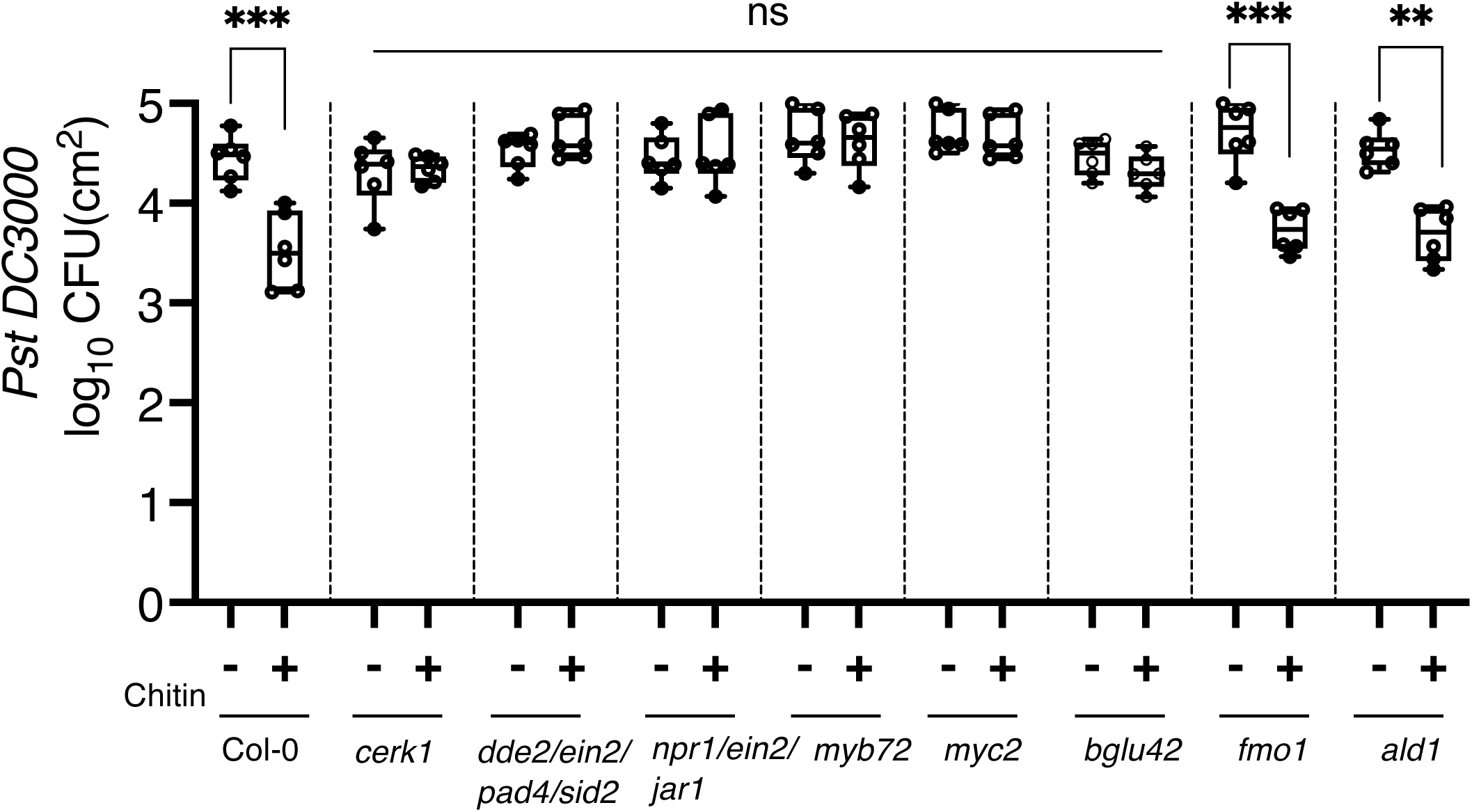
Chitin-induced systemic effects in Arabidopsis are mediated by ISR, but not SAR pathway. Four-week-old *Arabidopsis thaliana* Col-0 and mutants of ISR (*dde2/ein2/pad4/sid2*, *npr1/ein2/jar1*, *myc2*, myb72, *bglu42*) or SAR (*fmo1*, *ald1*) grown in chitin (+) vs non-chitin (-) amended soil were syringe-infiltrated with *Pst* DC3000 at a concentration of OD_600_=0.0002. Harvested leaf disks were ground and plated for colony counting. Values are mean ± SE, (n=6). Asterisks indicate a significant difference based on the Student’s *t*-test (**, P≤0.01; ***, P≤0.001).

### Chitin systemic effects on dieases resistance are independent of microbes from soil or chitin flakes

Given the proposed role of microbes in the ISR pathway (Vlot et al., 2021), we aimed to evaluate how soil microorganisms influence the systemic effects of chitin. We sterilised the soil and chitin flakes to eliminate microbial interactions, thereby isolating the effects of chitin itself (Lees et al., 2018; Williams-Linera & Ewel et al., 1984). The observed systemic effects were directly attributed to the application of chitin itself, independent of microorganisms present in soil or on chitin flakes (Fig. S6).

While it is known that microbes can play a role in ISR, our findings suggest that chitin is directly perceived by the plant, bypassing the need for microbial-mediated processes. One possible explanation is that upon detecting a small number of chitin oligomers present in crab chitin flakes, plant roots emit chitinases into the rhizosphere, potentially speeding up the chitin degradation process, even without the presence of soil microbes (Gianfreda, 2015; Tesfaye et al., 2005; Wasaki et al., 2005). However, we cannot rule out the possibility that microbes might amplify ISR through yet-unknown mechanisms. Our findings reveal that chitin directly enhances plant defense independent of soil microbes via direct perception in the roots broadening our understanding of its impact on systemic effects in distance tissues.

### Chitin soil amendment leads to systemic trancriptional potentiation of PTI components

PTI components, including PRRs/co-receptors, play a crucial role in mediating PAMP/MAMP-triggered immune responses. BIK1 acts as a signaling hub immediately downstream of PRRs and is involved in phosphorylation of downstream substrates, such as the NADPH oxidase RBOHD responsible of apoplastic production of ROS, whilst BAK1, serves as the primary co-receptor for various PRRs, enabling signal transduction (DeFalco & Zipfel, 2021). FLS2 acts as the main receptor for the bacterial MAMP/PAMP flg22 derived from bacterial flagellin, (Gómez-Gómez,& Boller, 2000). Markedly, upon chitin soil amendment, we observed an increase in protein levels of BIK1 and RBOHD (Fig. 7A), while FLS2 and BAK1 remained unchanged. Notably, these changes in protein abundance were linked to an increase in their transcript levels (Fig. 7B). These findings suggest that chitin-induced systemic effects involve specific changes in protein abundance of key PTI signaling components likely via transcriptional activation or post-translational regulation.

**Figure 7:**
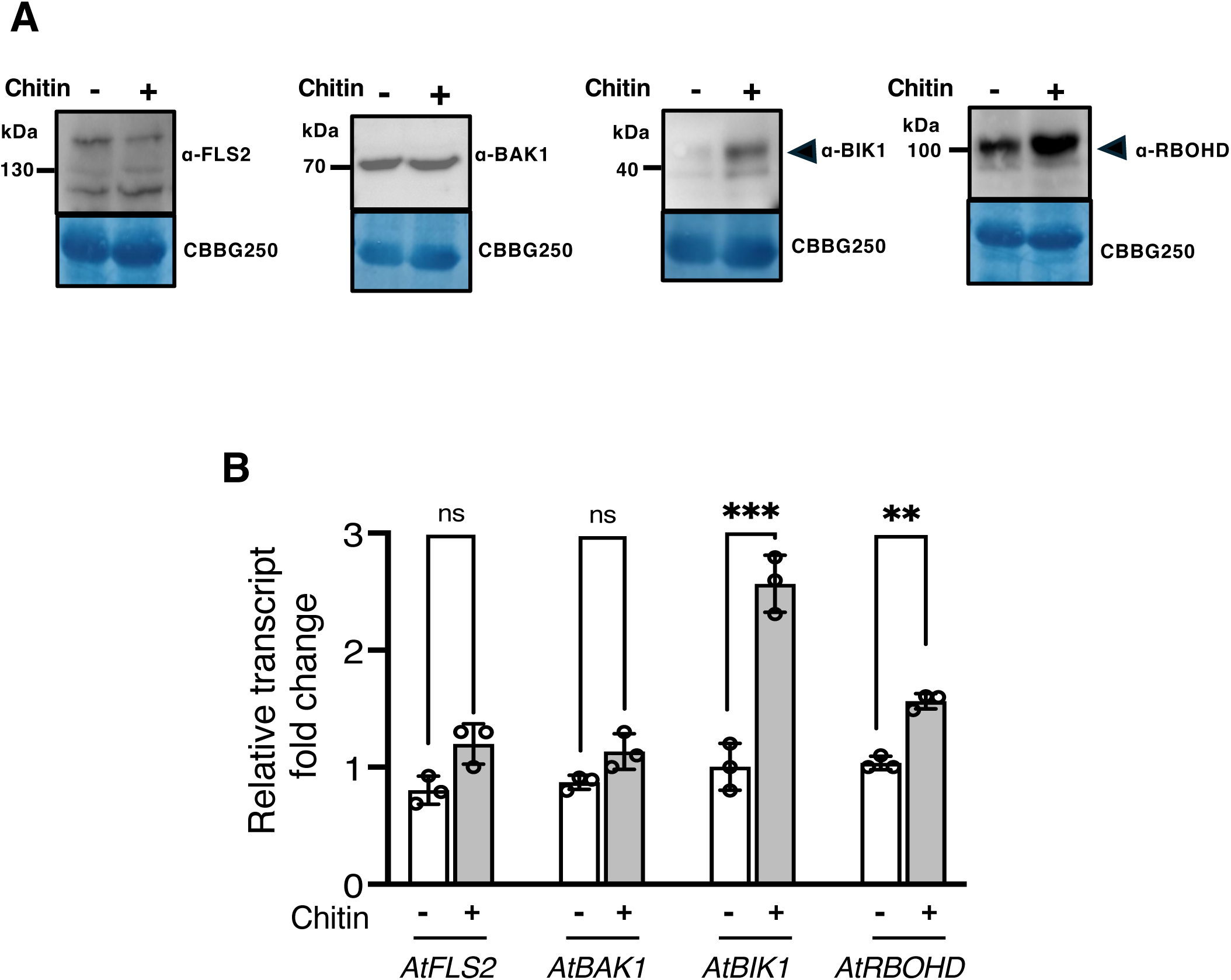
*BIK1* and *RBOHD* are transcriptionally upregulated upon chitin soil amendment. (A) Protein accumulation of PTI-related components after chitin soil amendment. Leaf samples were collected from 4-week-old Col-0 plants grown in soil amended (+) or not (-) with chitin (B) RT-qPCR analysis of PTI components (*BAK1*, *BIK1*, *FLS2*, *RBOHD*) expression levels. The same samples in (A) were used for RT-qPCR. *ACTIN* was used as a reference housekeeping gene. Values are mean ± SE, (n=3). Asterisks indicate a significant difference based on Student’s t-test (**, P≤0.01; ***, P≤0.001).

## Conclusion

In summary, this study provides evidence for the potential of chitin as a soil amendment biostimulant that enhances PTI and plant immune defenses against pathogens in leaves, detailing the molecular mechanisms involved. We demonstrated that chitin perception in roots enhances PTI in leaves across multiple plant species. Further analysis confirmed the essential role of active chitin perception in roots for mediating systemic effects, including systemic disease resistance, likely via the ISR pathway but independently of soil microbes. Changes in protein and transcript abundance of key PTI components shed light on the molecular mechanisms driving these systemic responses that are likely based on PTI potentiation. The nature of the systemic signal(s) and transcriptional mechanisms involved remain however to be deciphered. In any case, our results further reinforce the potential use of chitin in agricultural practices to boost crop disease resistance.

## Experimental procedures

### Plant material and growth conditions

*Arabidopsis thaliana* (Arabidopsis) seeds were subjected to surface sterilization, followed by stratification for three to five days at 4 °C. The sterilized seeds were then planted on Peat-based potting soil (‘einheitserde special’) and allowed to grow up to three to four true leaf stage before being transplanted into chitin and non-chitin-amended soil. Plants were kept in a growth chamber with controlled conditions of 20 °C temperature, 65 % humidity, and a 14-h light/10-h dark photoperiod. After the growth period, induced resistance assays were conducted.

Lines used in this study include Col-0 used as control, *fls2* (SAIL_691_C04; Zipfel et al., 2004); *cerk1* (Rosso et al., 2003); *fls2/efr-1/cerk1* (*fec*) (Xin et al., 2016); *lore* (SAIL_857_E06) (Ranf et al., 2015); *bak1-5* (Schwessinger et al., 2011); *bak1-5/bkk1* (Roux et al., 2011); *lym1/3* (Willmann et al., 2011); *fmo1* (Alonso et al., 2003); *ald1* (Song et al., 2004); *myc2* (Boter et al., 2004); *bglu42* (Alonso et al., 2003); *myb72* (Alonso et al., 2003); *ein2-1/jar1-1/npr1-1* (NASC ID no., N3801); *dde2-2/ein2-1/pad4-1/sid2-2* (Tsuda et al.,2009). Aequorin lines of Arabidopsis have been described previously (Knight et al., 1991) and were used for calcium influx measurements.

For *Lactuca sativa* L. (lettuce) variety Salinas, the trays were filled with peat-based sowing soil (N:P:K, 14:16:18) (Patzer Erden, Sinntal-altengronau). Subsequently, one seed was carefully sown using tweezers in the centre of each well and covered with a thin layer of sowing soil. Seedlings were transplanted into 18 well pots filled with potting soil containing chitin or not, at the three to four true leaf stage. Plants were kept in the greenhouse for at least seven days after being transferred in the controlled conditions of 21 °C and 16-h light/8-h dark photoperiod.

For *Triticum aestivum* (wheat) 20 °C/ 16 °C, 60 % relative humidity, 16/8 h photoperiod with a light intensity of 300 μmol/(s·m2) was used. JIW2-susceptible wheat cultivar Chinese Spring was grown in a growth chamber for 10-12 days in chitin and non-chitin mixed pots filled with potting soil. Detached segments of the first leaves were collected and inoculated with the *Bg. Tritici* isolate JIW2. Following incubation at 20 °C, 16 h light/ 8 h dark, and 80 % relative humidity, the experiment was scored 9–10 days after inoculation using a 1 to 100 % susceptibility scale, i.e. the leaf area covered with mildew was ranked phenotypically where lines with 100 % leaf area covered with mildew were considered fully susceptible and 0 % marked complete resistance (Kaur et al.,2008).

For *Solanum lycopersicum* (tomato), the same soil and procedure as for lettuce was used with slight modifications on the controlled growth conditions set at 20 °C and 16-h light/8-h dark photoperiod. Tomato Rio Grande WT and Bti9 are from Zeng et al., (2012).

### Chitin treatment

Crab shell-derived chitin flakes (BioLog Heppe GmbH, Lot: 40201609, Landsberg, Germany) were used in the study. Peat-based potting soil (Einheitserde special) was purchased from Patzer Erden, Sinntal-altengronau (Germany). The control soil consisted of potting soil without chitin addition, while the chitin-amended soil was prepared by mixing 2 g.L^-1^ of chitin with the potting soil. Both soil types were slightly moistened with water and subsequently incubated in sealed bags in the greenhouse for one week before use in the experiments.

### Bacterial infection assays

Bacterial infection assays were conducted following the protocol described by Zipfel et al., (2004) with slight modifications. Overnight cultures of *Pseudomonas syringae* pv. tomato (*Pst*) DC3000 were prepared in King’s B medium supplemented with 10 μg.mL^-1^ rifampicin and incubated at 28 °C with shaking at 220 rpm. For induced resistance assays, four-to five-week-old Arabidopsis, tomato and lettuce plants that had been grown with or without chitin were used. Three leaves from each plant were infiltrated with *Pst* DC3000 at OD_600_=0.0002 or approximately 1×10^5^ colony-forming units (CFU)/mL. Before inoculation, the overnight bacterial culture was subcultured in fresh LB medium for 1-2 h. The refreshed bacterial cells were harvested by centrifugation and resuspended in 10 mM MgCl2. After 48 h of inoculation, two leaf discs were collected from each infiltrated leaf, ground in 10 mM MgCl2, and plated on LB medium supplemented with 10 μg.mL^-1^ rifampicin. Plates were incubated at 28 °C for two days, and bacterial growth was quantified by counting CFUs.

### Blumeria graminis f. sp. tritici inoculation assay

The powdery mildew disease causing *B.g tritici* JIW2 isolate was used, given the wheat cultivar Chinese Spring’s susceptibility to this specific strain. The growth and inoculation procedures for wheat plants were based on Sánchez-Martín et al. (2016) with slight modifications. Briefly, the first leaves of 10-to 12-day-old wheat plants were cut to 2-3 cm long. Plastic plates were used as guides, and the cut leaf segments were immediately placed with their adaxial side facing up and abaxial side touching the agar plates, supplemented with 30 ppm benzimidazole. The infection chamber was prepared by ensuring cleanliness and placing the infection tower on top of opened plates. Infected leaf samples were obtained by taking leaf pieces that were already infected with powdery mildew. The infected leaf pieces were gently tapped on a folded piece of paper to dislodge the powdery mildew spores, which were then collected using a Pasteur pipette. Once the spores were collected, the spores were blown into the tower using the pipette. This process facilitated the introduction of powdery mildew spores into the infection tower, allowing them to settle on the wheat leaf segments. For each plate, equal amount of spores wereused. Disease levels were evaluated as per Kaur et al., 2008, 7-9 days after inoculation and incubation at 20°C, 16 h light/ 8 h dark, 80 % relative humidity.

### Reactive oxygen species (ROS) production assay

ROS production assays were conducted following the methodology described by Kadota et al., (2014) with minor modifications. In brief, 4-mm leaf disks were sampled from Arabidopsis, lettuce, or tomato plants. At least twelve leaf disks per genotype and condition were collected from 4-to 5-week-old plants. The leaf disks were placed in a 96-well plate filled with water and incubated overnight. On the following day, the water in the wells was replaced with a solution containing 1 μM L-012, 20 μg.mL^-1^ horseradish peroxidase (Sigma-Cat#77332), and 100 nM flg22 peptide (QRLSTGSRINSAKDDAAGLQIA) (SciLight Biotechnology LLC) or elf18 peptide (Ac-SKEKFERTKPHVNVGTIG) (EZBiolabs). Luminescence was measured using a Tecan SPARK multimodal plate reader (Tecan, Männedorf, Switzerland) for 42 min after flg22/elf18 or mock treatment.

### Calcium measurements

Leaf disks of Arabidopsis Col-0 carrying an aequorin transgene under the control of the 35S promoter (Knight et al., 1991) were harvested from 4-to 5-week-old plants using a 4-mm diameter biopsy punch and transferred into white 96-well plates. Each well contained 100 μL water. The leaf disks were allowed to rest overnight in 100 μL of 20 μM coelenterazine solution in the dark. The following morning, the coelenterazine solution was replaced with 100 μL of water, and the leaf disks were rested for at least 30 min in the dark. Readings were taken using a Tecan SPARK multimodal plate reader (Tecan, Männedorf, Switzerland) before and after adding 100 μl of 100 nM concentrated elicitor solution (flg22) or a mock solution. For each well, the readings were normalized to the average relative light unit (RLU) value obtained before the addition of the elicitor solution (L0). The total reconstituted aequorin was assessed by measuring the discharged aequorin with an excess of Ca^2+^ according to the method described by Knight et al., (1996, 2004). This resulted in luminescence proportional to the Ca^2+^ level, represented as the L/Lmax ratio. Here, L referred to the luminescence of aequorin at any given measurement, normalized to Lmax, which was the maximum luminescence remaining in the sample.

### Protein extraction and western blot

Protein extraction was done using the method previously described by Dindas et al., (2022). The membrane was blocked and probed in TBST-5 % non-fat milk by incubating with specific antibodies, including anti-BAK1 at 1:10000 dilution (Roux et al., 2011), anti-BIK1 at 1:5000 dilution (Agrisera-Cat#AS16 4030), anti-FLS2 at 1:1000 dilution (Chinchilla et al., 2006), anti-RBOHD at 1:3000 dilution (Agrisera - Cat#AS15 2962). The western blots were imaged using a Bio-Rad Chemidoc imaging system (Bio-Rad Laboratories, California, USA). To confirm equal loading of protein, the blotted membrane was stained with Coomassie brilliant blue (Abcam-Cat#ab119211).

### RNA extraction and real-time quantitative PCR (RT-qPCR) analysis

Total RNA was isolated from the target tissues using the FavorPrep Plant Total RNA Purification Mini Kit (Favorgen) following the manufacturer’s instructions. First-strand cDNA synthesis was performed using 1 μg of DNA-free RNA sample with the RevertAid First Strand cDNA Synthesis Kit (Thermo - Cat#EP0441) according to the manufacturer’s protocol. For RT-qPCR analysis, the diluted cDNA was used as the template with PowerUp SYBR Green (Thermo-Cat#A25741), and the specific primers listed in Table S1. Data analysis was carried out using the ΔΔCT method, and the expression levels were normalized to the AtACTIN reference gene. Three biological replicates, each consisting of three technical replicates, were performed for each sample.

### Micro-grafting of Arabidopsis

The micrografting technique was employed to create chimeric Arabidopsis plants, following a modified version of the protocols described by Vanderstraeten et al., (2022) and Tsutsui et al., 2020. The process began by surface-sterilizing Arabidopsis seeds, which were then plated on agar-containing Murashige and Skoog (MS) media supplemented with vitamins, 1 % sucrose, and 0.8 % agar, stratified for 2-3 days in the dark at 4 °C. After a period of stratification, the plates were transferred to vertical growth conditions to induce hypocotyl etiolation. Grafting was performed under sterile conditions, with the hypocotyls of young Arabidopsis (Col-0 WT and *cerk1* mutants) seedlings carefully cut and scions and rootstocks joined and aligned on a new plate. The plants were allowed to heal and grow under standard conditions, and grafting efficiency was evaluated based on the success of the union between scion and rootstock.

### Phylogenetic analysis

Protein sequences for analysis were sourced from the lettuce genome database, using BLAST of conserved kinase domain sequences of the Arabidopsis LysM receptor kinase family (LYKs), which include AtCERK1 (At3g21630); AtLYK2 (At3g01840); AtLYK3 (At1g51940); AtLYK4 (At2g23770) and AtLYK5 (At2g33580) to find AtLYKs protein homologues at the UC Davis lettuce genome repository (https://lgr.genomecenter.ucdavis.edu) (Reyes-Chin-Wo et al., 2017), yielding the LsLYK sequences (Fig. S1). The alignment of these sequences was executed using the MUSCLE algorithm within the MEGA X software (Kumar et al., 2018). For phylogenetic analysis, we constructed rooted trees using MEGA X, utilizing the UPGMA algorithm under default settings and performed bootstrap validation with 1000 iterations. The resulting phylogenetic trees were visualized and interpreted using iTOL (https://itol.embl.de/) (Letunic and Bork, 2019).

### Plasmid construction and generation of transgenic plants

The *AtCERK1* homologue in lettuce *LOC111905667* (*LsCERK1*) was targeted to generate *Lscerk1* mutants in lettuce using CRISPR-Cas9-mediated mutagenesis. Cloning of *LsCERK1* level 0 and level 1 parts into the PiCSL32281+LacZ vector was facilitated using the Golden Gate modular cloning system, as per Castel et al. (2019). The RUBY selection marker (He et al., 2020) was under the control of the 35S promoter, fused to RPS5-driven Cas9. Each cassette was assembled into level 1 via Bsa1 enzyme (5’ -GGTCTC-3’) before being assembled into the final level 2 destination vector p35S::RUBY::Cas9::gRNAs::EL (Fig. S2) via BpiI enzyme (5’ -GAAGAC-3’). Two guide RNAs were designed: one at the 5’ end of the first intronless exon of *LsCERK1* and another one at 130 nt downstream.

The transformation into lettuce was achieved through modifications of the protocol of Bertier et al. (2018). In brief, *Agrobacterium tumefaciens* strain GV3101 was used to transform 3-to 4-day-old Salinas lettuce cotyledons. This was followed by a co-cultivation period and callus induction, critical for successful genome integration. The callus was transferred into shoot and root-forming media. Successfully transformed lettuce plantlets were easily identified by their expression of the *RUBY* gene. To verify the genetic constructs and mutations, we employed Sanger sequencing (Fig. S3). The *Lscerk1* line has a 128-bp deletion in the first exon of the *LsCERK1* gene, causing a frameshift (Fig. S3).

### Data analysis

Statistical analysis was carried out using the GraphPad Prism 9.0 software. The significance of differences between groups was determined based on a threshold of P<0.05, indicating that differences with a probability of less than 5 % were considered statistically significant.

## Acknowledgements

We thank Greg Martin, Jürgen Zeier, Thorsten Nürnberger, and Zachary Nimchuk for generously providing published plant lines and protocols. We thank Hiroki Tsutsui for his advice on micrografting techniques; Ashley Crook, Matthew Moscow, and David Tricoli for their help with lettuce transformation techniques, and the members of the Zipfel laboratory and ChitinOMix consortium for useful discussions. This project was funded by grant no. 189340 from the Swiss National Science Foundation (SNSF) and grant no. G063820N from the Fonds Wetenschappelijk Onderzoek (FWO) for the consortium ChitinOMix. Research in the CZ and BK laboratories is also funded by the University of Zurich. YG was partially supported by a Swiss Government Excellence Scholarship, an EMBO Postdoctoral Fellowship (ALTF 386-2021) and a JSPS Overseas Research Fellowship.

## Author contributions

MM, YG, and CZ conceived and designed the experiments; MM, HZ, and VW conducted the experiments and MM analyzed the data. MM wrote the first draft of the manuscript, with input from all authors. CZ, BK, and YG supervised the project and contributed to the writing. CZ and BK obtained funding.

## Supporting Information

For further supplementary details, please refer to the Supporting Information section located after the article.

**Figure S1: Phylogenetic analysis of the LsLYK family.** BLAST searches were performed using the conserved kinase domain sequences of Arabidopsis LYKs (CERK1/LIK1, LYK2, LYK3, LYK4 and LYK5) to lettuce genome database. The multiple alignment was executed using the MUSCLE algorithm within the MEGA X software. The rooted tree using FERONIA (AT3G51550) as an outgroup was constructed using MEGA X constructed using MEGA X, utilizing the UPGMA algorithm under default settings and performed bootstrap validation with 1000 iterations. The phylogenetic tree was visualized and interpreted using iTOL. In bold are AtLYKs family (in black) and their homologues in lettuce (in red).

**Figure S2: Golden Gate CRISPR Level 2 construct used for *Lscerk1* mutant generation.** This schematic depicts (a) the *LsCERK1* gene map with 12 exons and 2 gRNAs target (blue down arrows) for generating *Lscerk1* mutant with a 128-nt deletion. The corresponding LsCERK1 domains contain the signal peptide (lavender colour), LYM domain (gray colour), Transmembrane domain (TD) (light grey colour), and kinase domain (KD) (rose-brown colour). The positions of the two gRNAs used and genotyping primers are shown in light green and purple colour respectively. (b) Depicts the assembled Golden Gate system used for CRISPR-Cas9 Level 2 construct (Castel et al., 2019) aimed at the *LsCERK1* gene in lettuce. Two guide RNAs were designed, one at the 5’ end of the first intronless exon of LsCERK1 and another one at 130 nt downstream. RUBY was used as visible selection marker (He et al., 2020) . Each cassette was assembled into level 1 via Bsa1 enzyme (5′-GGTCTC-3′) before being assembled together into the final destination vector level 2 via BpiI enzyme (5′-GAAGAC-3′).

**Figure S3: *Lscerk1* mutant chromatograms of the LYM domain sequencing.** Sequences of *Lscerk1* mutants used to characterize the mutations in the *Lscerk1* gene showing alignments of the *LsCERK1.* Reference (Ref) amino acid sequences alongside WT Salinas and *Lscerk1*, which exhibits a 128-bp deletion in the Lysin Motif (LYM) domain. This deletion leads to a frameshift and introduces early stop codons (red stars). The regions targeted by gRNA1 and gRNA2 within the LYM domain are marked by blue downward arrows. Chromatograms from Sanger sequencing illustrate the sequence of WT alongside the *Lscerk1* mutant, highlighting the absence of 128 bp in the *Lscerk1* due to the deletion.

**Figure S4: Chitin-triggered ROS production is abolished in *Lscerk1* mutant.** Leaf disks from 4-week-old *Lactuca sativa* L cv. Salinas WT and *Lscerk1* mutant plants were treated with 50 ug/mL of chitin polysaccharide and mock. Comparative cumulative ROS production in WT and *Lscerk1* mutant lines after chitin treatment shows the diminished ROS production in the mutants. Values are mean ± SE, (n=12). Different characters indicate significant differences based on one-way ANOVA and Tukey’s post hoc test (P≤0.05).

**Figure S5: Flg22-induced ROS production potentiation is compromised in ISR mutants.** Leaf disks from 4-week-old *Arabidopsis* thaliana plants Col-0, mutants of ISR (*dde2/ein2/pad4/sid2*, *npr1/ein2/jar1*) and SAR (*fmo1*, *ald1*) all grown in chitin (+) vs non-chitin (-) amended soil were treated with 100 nM flg22. Total ROS RLU was quantified 42 min post-treatment. Values are mean ± SE, (n=12). Asterisks indicate a significant difference based on Student’s ttest (*, P≤0.01).

**Figure S6: Chitin-induced systemic effects are independent of microbes from soil or chitin flakes.** Four-week-old *Arabidopsis thaliana* plants Col-0 WT and *cerk1* were grown in different combinations of either sterilised soil or sterilised chitin or both. (A) Unsterilised soil + sterilised chitin. (B) Unsterilised soil + Unsterilised chitin. (C) Sterilised soil + Unsterilised chitin and, (D) Sterilised soil + sterilised chitin. Plant were grown under the conditions of chitin (+) vs non-chitin (-) treated soils as well as sterilised and non sterilised soil combinations. Plants were syringe-infiltrated with *Pst* DC3000 at a concentration of OD_600_=0.0002. Harvested leaf disks were ground and plated for colony counting to quantify bacterial titers two days post-infiltration. Values are mean ± SE, (n=6). Asterisks indicate a significant difference based on Student’s t-test (*, P≤0.01).

**Figure S7: Working model for chitin-induced systemic disease resistance.** Upon pathogen attack, plants initially respond in a natural unprimed state (1). When chitin is added to the soil, it is recognized by chitin receptors AtLYK5 and AtCERK1 in roots. This recognition triggers a local response (2), while simultaneously initiating the priming process in distant tissues through the ISR pathway, facilitated by an unidentified mobile component (3). As a result, the plant becomes primed, leading to increased levels of PTI component proteins such as BIK1 and RBOHD (4). This priming is thought to confer a state of increased readiness in the plant, allowing for a stronger and more effective defense response upon subsequent pathogen attack. The observed suppression of pathogen growth in leaves is attributed to this enhanced defense response (5).

**Table S1:**
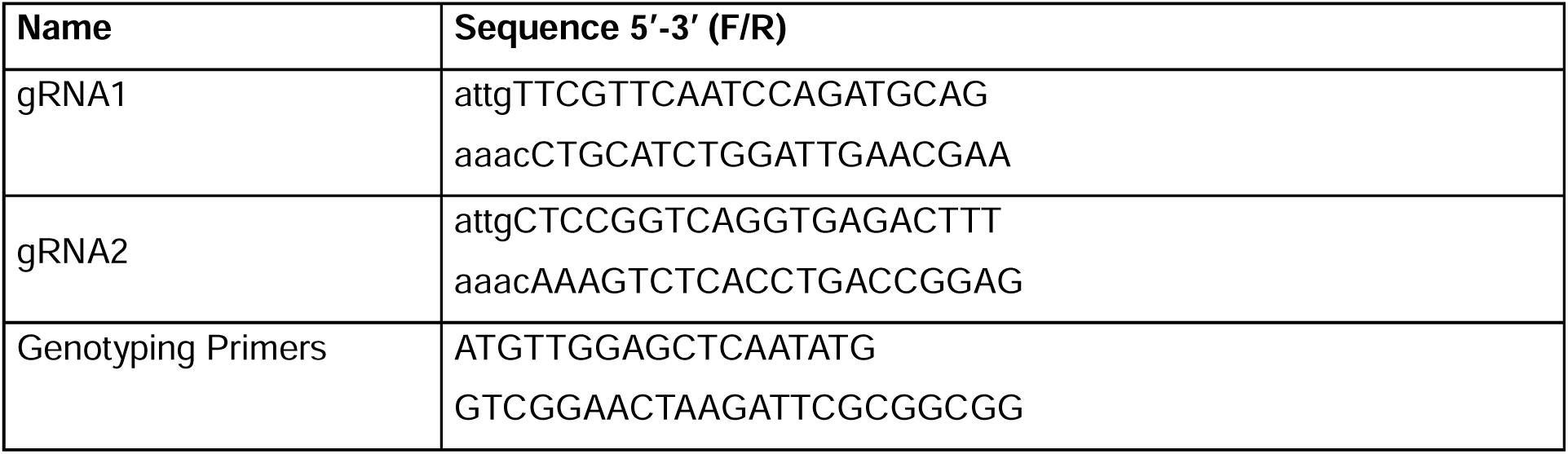
*LsCERK1* gRNAs sequences and genotyping primers used in this study.

**Table S2:**
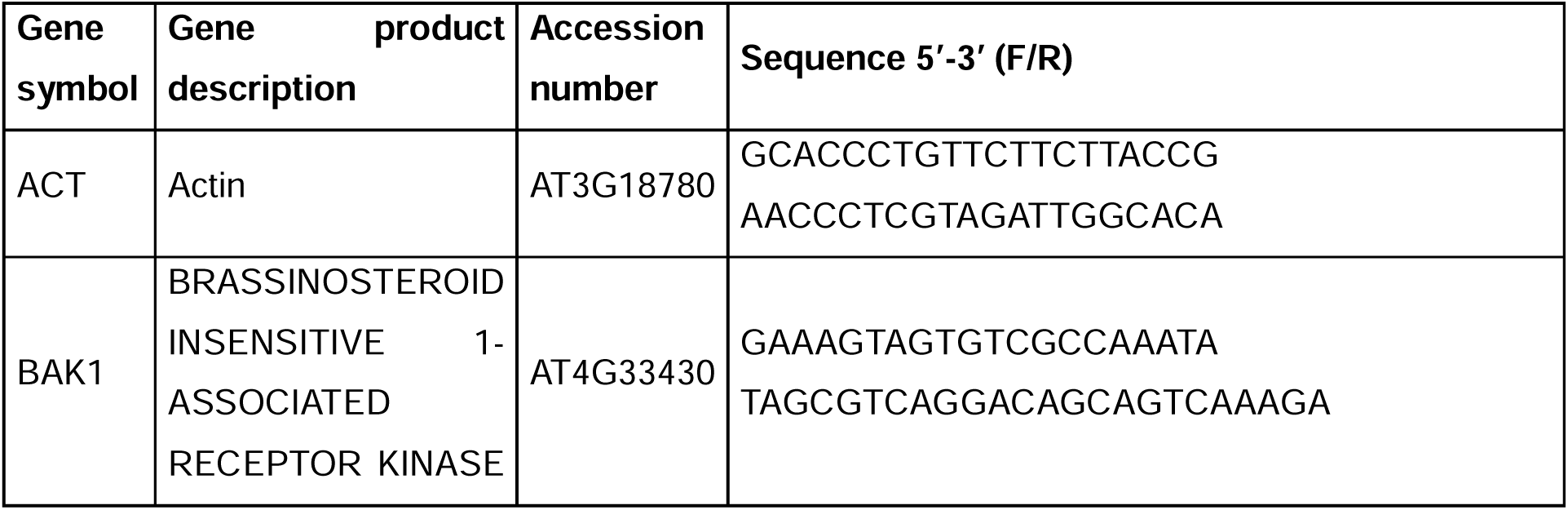
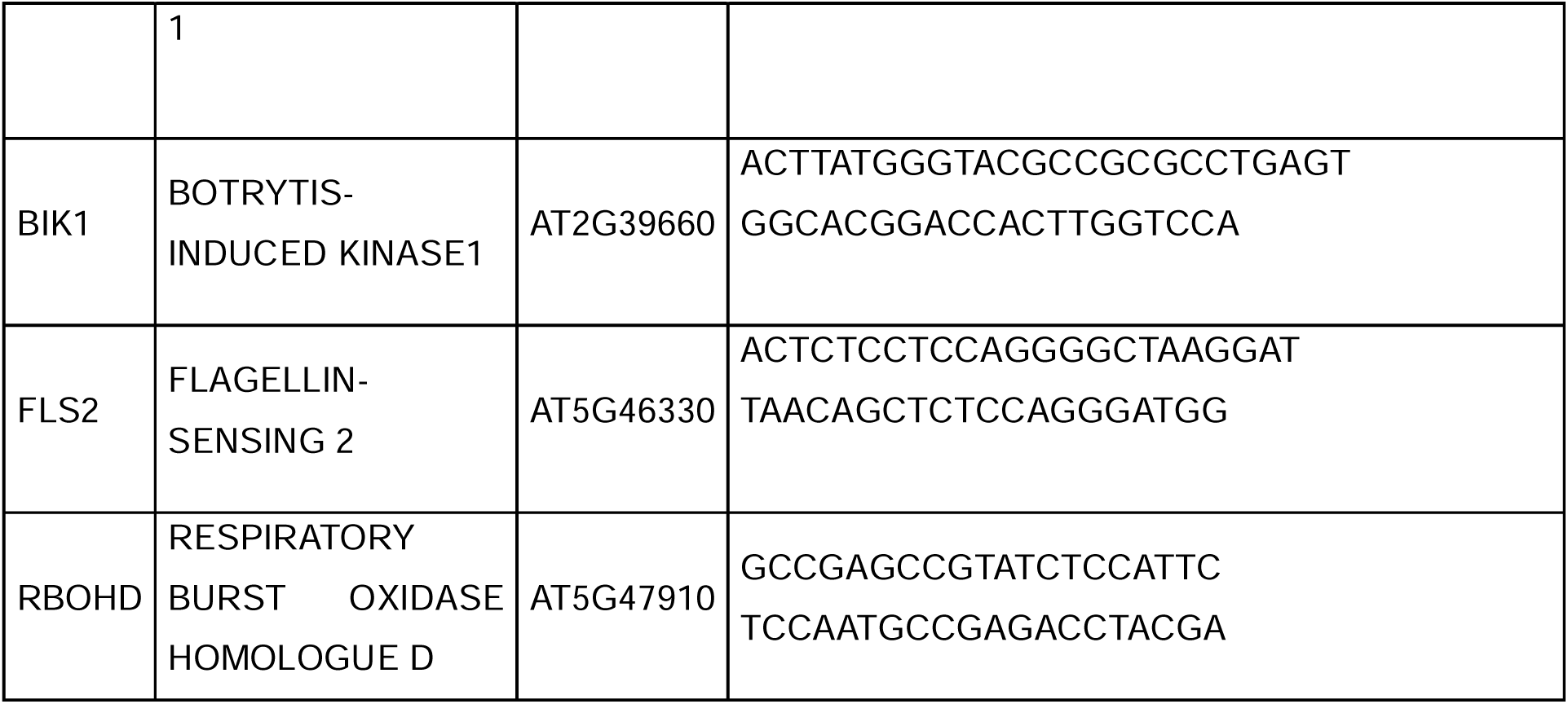
RT-qPCR primers used this study.

