## Supplementary Figures 1-7 for "Chitin soil amendment triggers systemic plant disease resistance through enhanced pattern-triggered immunity"

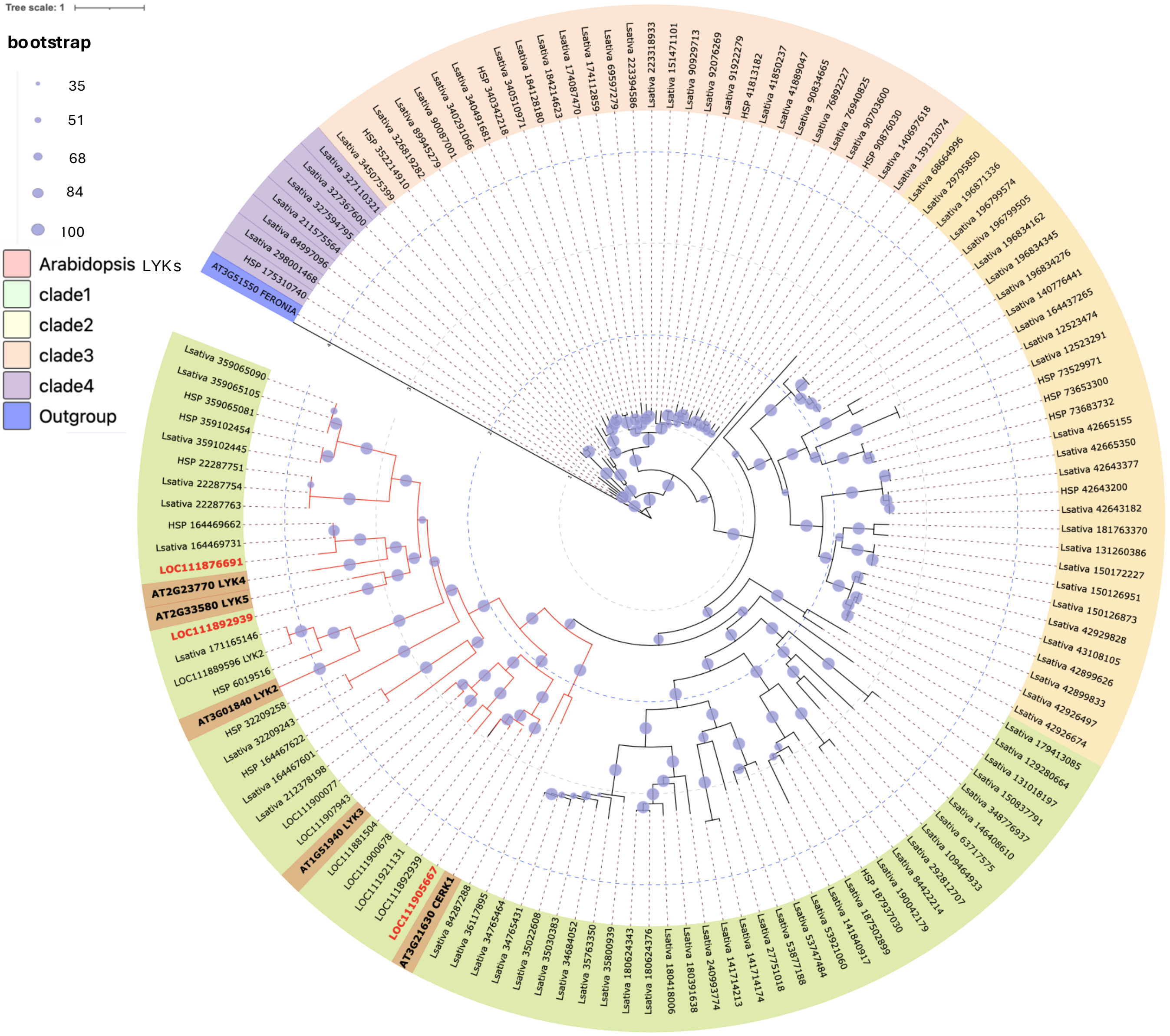


**Fig S1**: **Phylogenetic analysis of LsLYKs family**. BLAST of conserved kinase domain sequences of the *Arabidopsis* LYKs (CERK1, LYK2, LYK3, LYK4 and LYK5) family to lettuce genome database. The alignment was executed using the MUSCLE algorithm within the MEGA X software. The rooted tree using FERONIA (AT3G51550) as an outgroup was constructed using MEGA X, utilizing the UPGMA algorithm under default settings and performed bootstrap validation with 1000 iterations. The phylogenetic tree was visualized and interpreted using iTOL. In bold are AtLYKs family (in “black”) and their homologues in lettuce (in “red”).

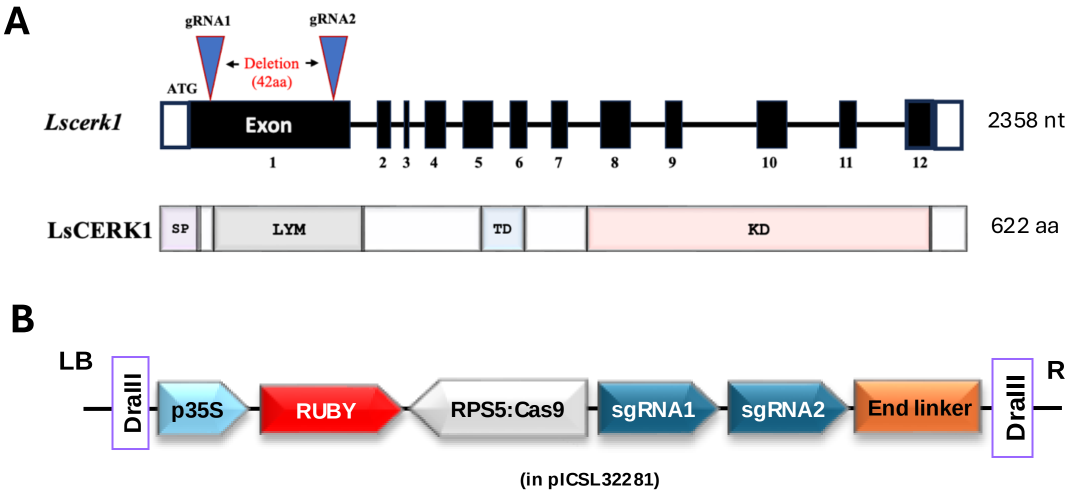


**Fig S2: *LsCERK1* gene map and the Golden Gate CRISPR Level 2 construct used for *Lscerk1* mutant generation.** This schematic depicts (a) the *LsCERK1* gene map with 12 exons and 2 gRNAs target (blue down arrows) for generating *Lscerk1* mutant with a 128 nt base deletion. The corresponding LsCERK1 domains contain the signal peptide (lavender colour), LYM domain (gray colour), Transmembrane domain (TD) (light grey colour ), and kinase domain (KD) (rose-brown colour). The positions of the two gRNAs used and genotyping primers are shown in light green and purple colour respectively. (b) Depicts the assembled Golden Gate system used for CRISPR-Cas9 Level 2 construct (Castel *et al.,* 2021) aimed at the LsCERK1 gene in lettuce. Two guide RNAs were designed, one at the 5' end of the first intronless exon of LsCERK1 and another one at 130 nucleotides downstream. RUBY was used as an easy -to- visualize selection marker (He *et al.,* 2020) . Each cassette was assembled into level 1 via Bsa1 enzyme (5′-GGTCTC-3′) before being assembled together into the final destination vector level 2 via BpiI enzyme (5′-GAAGAC-3′).

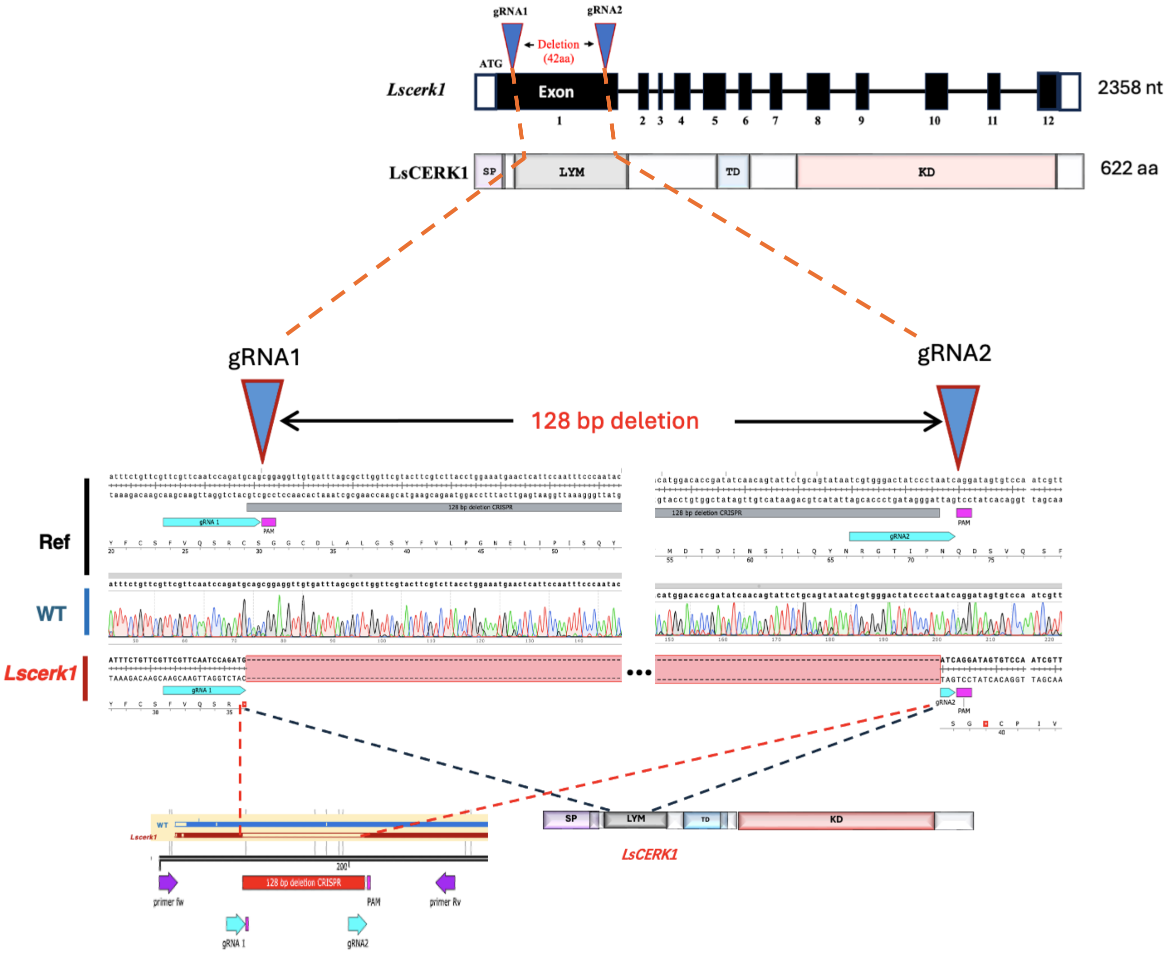


**Fig S3: *Lscerk1* mutant chromatograms of the LYM domain sequencing**.

Sequences of *Lscerk1* mutants used to characterize the mutations in the *Lscerk1* gene showing alignments of the LsCERK1 Reference (Ref) amino acid sequences alongside WT Salinas and *Lscerk1*, which exhibits a 128 bp deletion in the Lysin Motif (LYM) domain. This deletion leads to a frameshift and introduces early stop codons (red stars). The regions targeted by gRNA1 and gRNA2 within the LYM domain are marked by blue downward arrows. Chromatograms from Sanger sequencing illustrate the sequence of WT alongside the *Lscerk1* mutant, highlighting the absence of 128 bases in the *Lscerk1* due to the deletion.

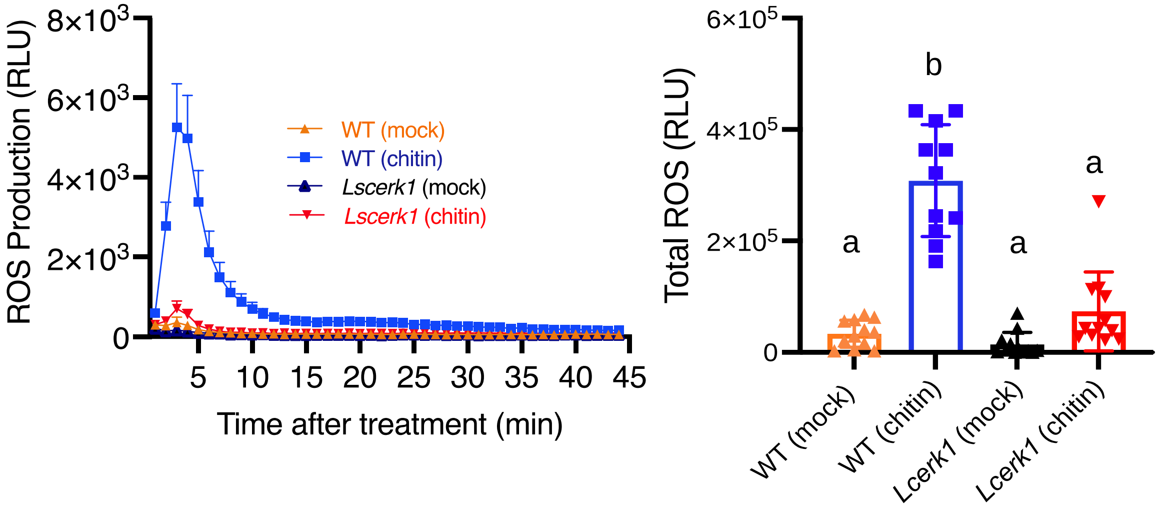


**Fig S4**: **Chitin-triggered ROS in *Lscerk1* mutants is abolished.** Leaf disks from 4-week-old *Lactuca sativa L cv. Salinas* and *Lscerk1* mutant plants were treated with 50 ug/mL of chitin polysaccharide and mock. Comparative cumulative ROS production in wild-type and *Lscerk1* mutant lines after chitin treatment shows the diminished ROS response in the mutants. Values are mean ± SE, (*n*=12). Different characters indicate significant differences based on one-way ANOVA and Tukey's post hoc test (P≤0.05).

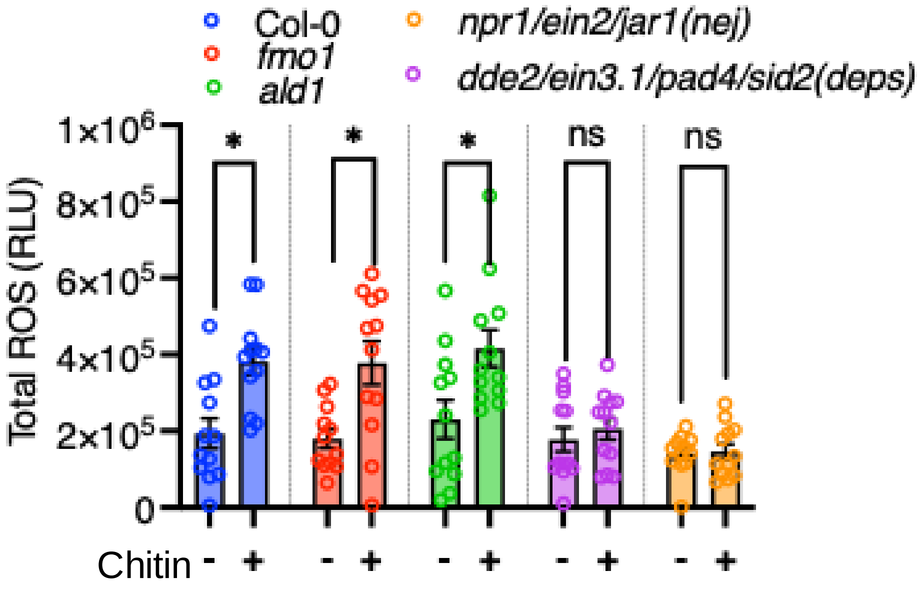


**Fig S5**: **flg22-induced ROS potentiation is compromised in ISR mutants:** Leaf disks from 4-week-old *Arabidopsis thaliana* plants Col-0; mutants of ISR (*dde2/ein2/pad4/sid2, npr1/ein2/jar1*) and SAR (*fmo1 & ald1*) all grown in chitin (+) vs non-chitin (-) were treated with 100 nM flgg22. Total ROS RLU was quantified 42 min post-treatment. Values are mean ± SE, (*n*=12). Asterisks indicate a significant difference based on Student's *t*-test (*, P≤0.01).

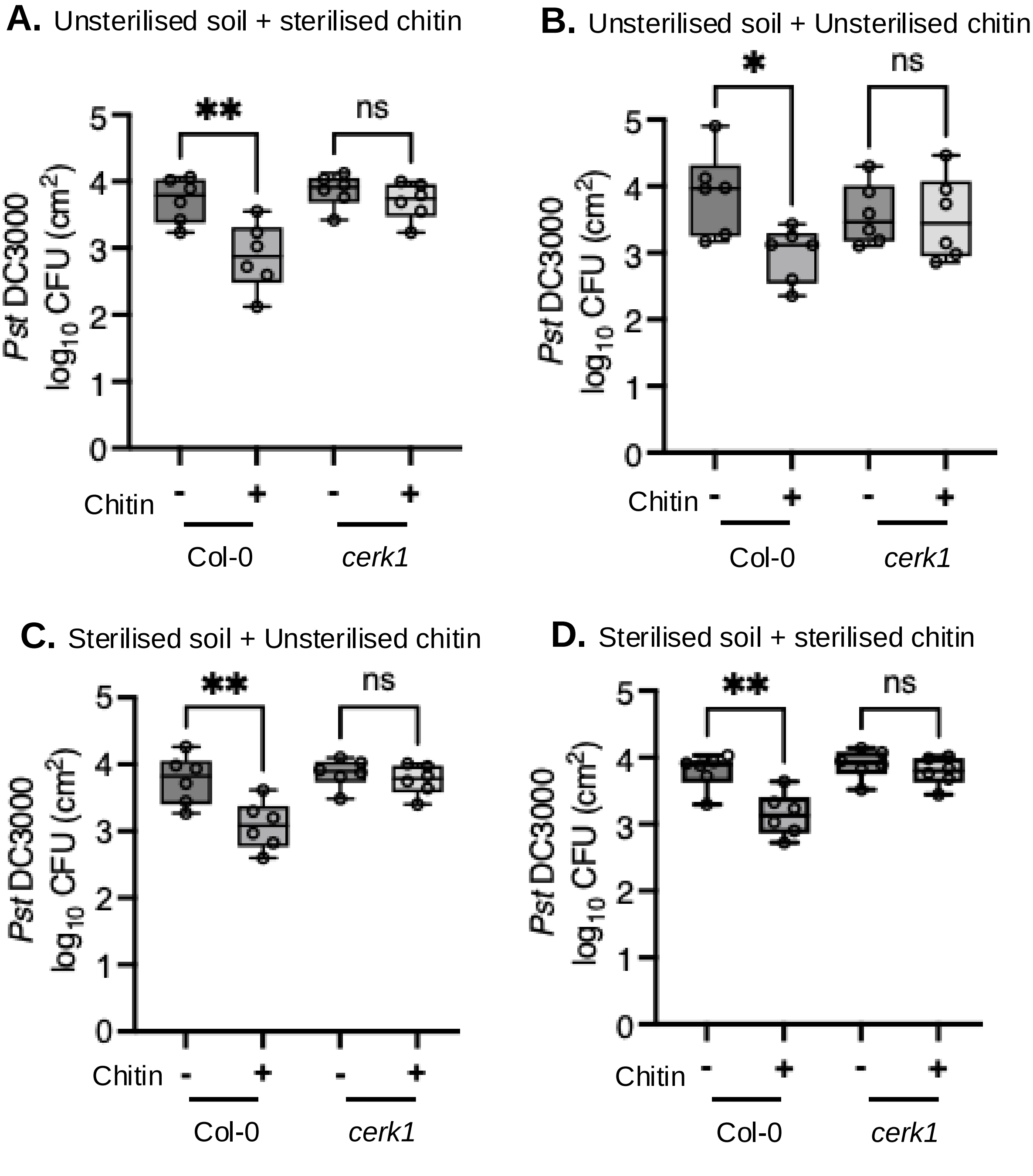


**Fig S6**: **Chitin-induced systemic effects are soil microbe independent.**
4-week-old *Arabidopsis thaliana* plants (WT Col-0 vs *cerk1*) were grown in different combinations of either sterilised soil or sterilised chitin or both. (A) Unsterilised soil + sterilised chitin. (B) Unsterilised soil + Unsterilised chitin. (C) Sterilised soil + Unsterilised chitin and, (D) Sterilised soil + sterilised chitin. The plants were allowed to grow under the conditions of chitin (+) vs non-chitin (-) treated soils of different soil and chitin sterilisation combinations. Plants were syringe infiltrated with *Pst DC3000* at a concentration of 0.0002 (OD600). Harvested leaf disks were ground and plated for colony counting to quantify pathogen growth 2 days post-infiltration. Values are mean ± SE, (*n*=6). Asterisks indicate a significant difference based on the Student's *t*-test (**, P≤0.01).

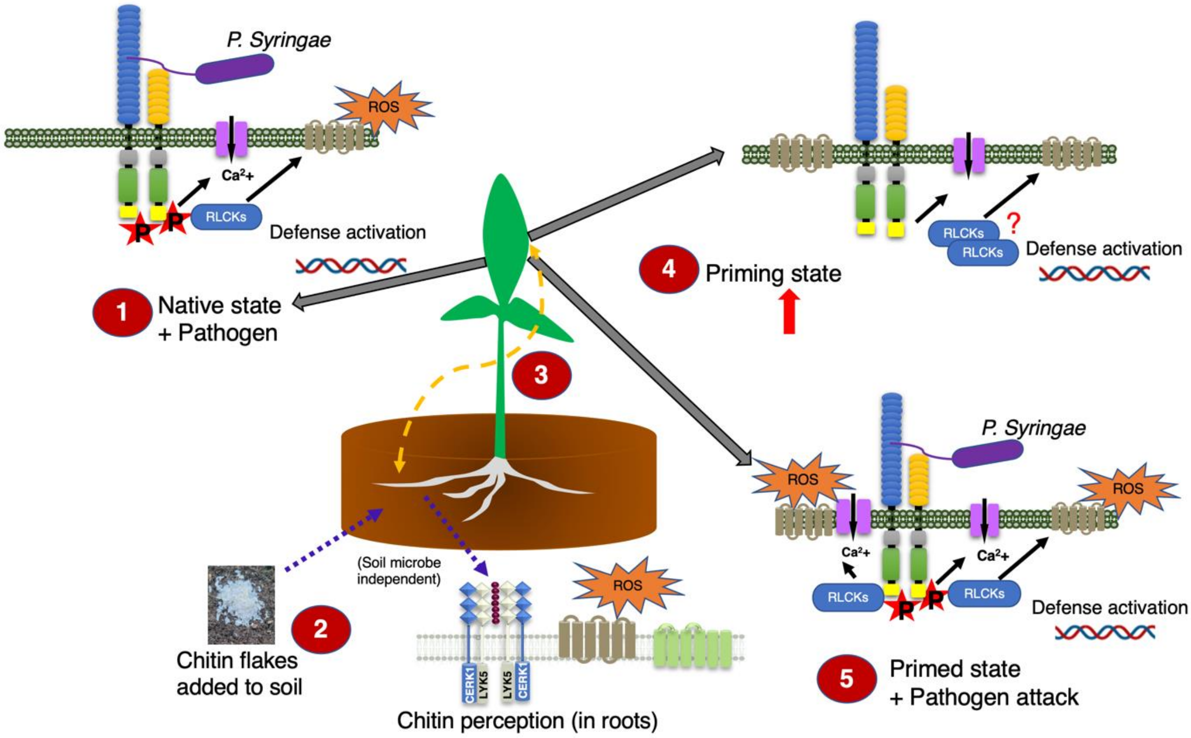


**Fig S7: Working model for chitin-induced systemic antipathogenic effects.** Upon pathogen attack, plants initially respond in a natural unprimed state **(1).** However, when chitin is added to the soil, it is recognized by chitin receptors, AtLYK5 and AtCERK1, present in the roots. This recognition triggers a local response **(2),** while simultaneously initiating the priming process in distant tissues through the ISR pathway, facilitated by an unidentified mobile component **(3).** As a result, the plant becomes primed, leading to increased levels of PTI component proteins such as BIK1 and RBOHD **(4).** This priming is thought to confer a state of increased readiness in the plant, allowing for a stronger and more effective defense response upon subsequent pathogen attack. The observed suppression of pathogen growth on the phyllosphere is attributed to this enhanced defense response **(5).**
